# Preclinical evaluation of a precision medicine approach to DNA vaccination in Type 1 diabetes

**DOI:** 10.1101/2021.06.02.446808

**Authors:** Jorge Postigo-Fernandez, Rémi J. Creusot

## Abstract

Antigen-specific immunotherapy involves the delivery of self-antigens as proteins or peptides (or using nucleic acids encoding them) to be presented with the goal of inducing tolerance. Approaches employing specific epitopes restricted to the subject’s MHC haplotypes have multiplied and offer a more focused and tailored way of targeting autoreactive T cells. In addition, the Endotope platform allows endogenously expressed epitopes to be processed and presented on appropriate MHC class I and II molecules. Here, we evaluated the efficacy of a DNA vaccine encoding epitopes selected and tailored for the non-obese diabetic (NOD) mouse compared to the expression of the proinsulin protein, one of the most successful antigens in prevention of NOD disease, and we assessed the influence of several parameters (e.g. route, dosing frequency) on preventing diabetes onset at normoglycemic and dysglycemic stages. First, encoded peptides should be secreted for effective disease prevention. Furthermore, short weekly treatments with Endotope and proinsulin DNA vaccines delay disease onset, but sustained treatments are required for long-term protection, which was more significant with intradermal delivery. Although epitopes can be presented for at least two weeks, reducing the frequency of antigen administration from weekly to every other week reduced efficacy. Finally, both Endotope and proinsulin DNA vaccines were effective in the dysglycemic stage of disease, but proinsulin provided better protection, particularly in subjects with slower progression of disease. Thus, our data support the possibility of applying a precision medicine approach based on tailored epitopes for the treatment of tissue-specific autoimmune diseases with DNA vaccines.

**SIGNIFICANCE STATEMENT:** Antigen-specific immunotherapy is a targeted approach to treat autoimmune diseases by turning off responses to disease-relevant antigens only, leaving the rest of the immune system unaffected. Protein antigens contain many epitopes, but only a fraction of them can be presented on a specific HLA haplotype and the relative importance of different antigens vary between patients due to disease heterogeneity. Strategies based on specific epitopes do not only consider the HLA haplotype and immune profile of groups of patients but can also include important neoepitopes not present in protein antigens. Here, we provide proof-of-principle that such strategy applied to tolerogenic DNA vaccination is effective in a preclinical model of autoimmune diabetes, paving the way for precision medicine using endogenously encoded epitopes.

## INTRODUCTION

Antigen-based approaches to treat autoimmune diseases have the attractive capacity to selectively block the disease-driving lymphocytes without impairing our overall immunity to pathogens and malignancies. There remains clear yet unmet clinical need for such therapies capable of effectively reestablishing tolerance to targeted autoantigens, but despite an excellent safety profile, they have not yet achieved this goal in patients.

Antigen-specific immunotherapies (ASITs) in Type 1 diabetes (T1D) are primarily targeted to diabetogenic CD4 and CD8 T cells that are reactive to multiple β-cell antigens (1, 2). The targeting is achieved by delivery and presentation of one or more of these antigens in a manner that results in deletion, regulation, anergy and/or exhaustion of these autoreactive T cells. Selected autoantigens have been administered to patients in the form of proteins or peptides via parenteral, oral or nasal routes, or in the form of protein-encoding DNA plasmids (DNA vaccines) (1, 3). A wide variety of delivery vehicles have been developed and tested to funnel these antigens to specific cell types and/or anatomical locations, including various micro- and nanoparticles (4), and more recently, soluble antigen arrays (5). A DNA-based approach allows the patient’s own cells to (i) endogenously express the protein(s) of interest, which may sometimes be challenging to produce recombinantly (e.g. proinsulin), (ii) apply natural post-translational modifications (PTMs), some of which are playing a critical role in the disease process (6, 7) and (iii) may allow the antigen(s) to persist longer than exogenous antigens (with the exception of nanoparticles designed to achieve slow antigen release).

Choosing the right antigen(s) to treat T1D is difficult(8). Historically, insulin/proinsulin and GAD65 have been favored due to successes in preclinical studies in non-obese diabetic (NOD) mice, the prevalence of anti-insulin and GAD65 autoantibodies and the identification of many T cell clones reactive to these antigens in T1D patients. In recent years, new and more complex islets autoantigens have emerged, including PTM versions and other neoantigens such as hybrid peptides (6, 9, 10).

At the same time, better understanding of the heterogeneity of T1D patients in disease progression, immune profile and genetic risk factors has led to their stratification into subgroups termed “endotypes” (11, 12). Another parameter considered in the stratification of patients is their responsiveness to particular types of therapies. In ASITs, responsiveness limited to a subset of patients in clinical trials (e.g. DPT-1 oral insulin (13, 14)) indicates that there is no “one-antigen-fits-all” option, and once responsiveness can be linked to endotype, it becomes possible to subsequently screen patients that are more likely to benefit from the treatment. Most recent clinical trials have expanded their eligibility criteria to include immune markers (e.g. presence of a specific autoantibody such as GADA in NCT02387164) or specific human leukocyte antigen (HLA) haplotypes (in the case of peptide-based therapy, for example HLA-DRB1*0401 in NCT02620332). Several studies suggest that presentation of pertinent epitopes with appropriate MHC restriction can circumvent the need to use full proteins, in part because of linked/bystander suppression (15-18). For specific HLA haplotype, several immunodominant or immunoprevalent epitopes have been identified across multiple antigens (9, 19-21).

We have designed constructs to achieve optimal presentation of endogenously expressed epitopes (Endotope platform) to CD4 and CD8 T cells (22). We previously reported that these constructs encoding both native epitopes and mimotopes could achieve engagement of the corresponding antigen-specific T cells in vitro (22) and in vivo (23) and significantly delay the onset of diabetes in NOD mice (23). Here, we assessed how multiple parameters influence the efficacy of Endotope plasmid DNA (pDNA) with epitopes tailored to the NOD mouse in preventing/delaying disease for preclinical optimization of this approach, including the localization of the expressed polypeptides, the route of inoculation, the duration and frequency of treatment, and the stage of disease at treatment initiation. In some instances, we compared its efficacy to the Proinsulin pDNA (24), used here as gold standard because it constitutes the basis of the current Phase II SUNRISE clinical trial (NCT03895437). In phase I studies, intramuscular inoculation of Proinsulin pDNA led to delayed loss of C-peptide, reduction of insulin-reactive CD8 T cells while showing an excellent safety profile (25).

## RESULTS

### Secretion of Endotope polypeptide is required for protection by a DNA vaccine

We previously reported that both AI and BS constructs (**Fig.1**) elicit antigen-specific T cell responses (albeit stronger with BS) in the draining lymph nodes after i.m. and i.d. treatment, and that delivery of a mix of AI and BS-containing pDNA significantly delayed diabetes onset in NOD mice after an 8-week i.m. treatment, similar to the Proinsulin DNA vaccine (23). To follow up on these studies, we evaluated constructs AI versus BS separately to ascertain their relative contribution to disease protection, using continuous treated via i.d. route, which we had not previously tested. The secreted form of Endotope appears to be the one mediating the protection from T1D, as the intracellular form had no effect on incidence (**Fig.2**). As a result, only the BS variant of Endotope was used in subsequent experiments.

**Fig. 1.**
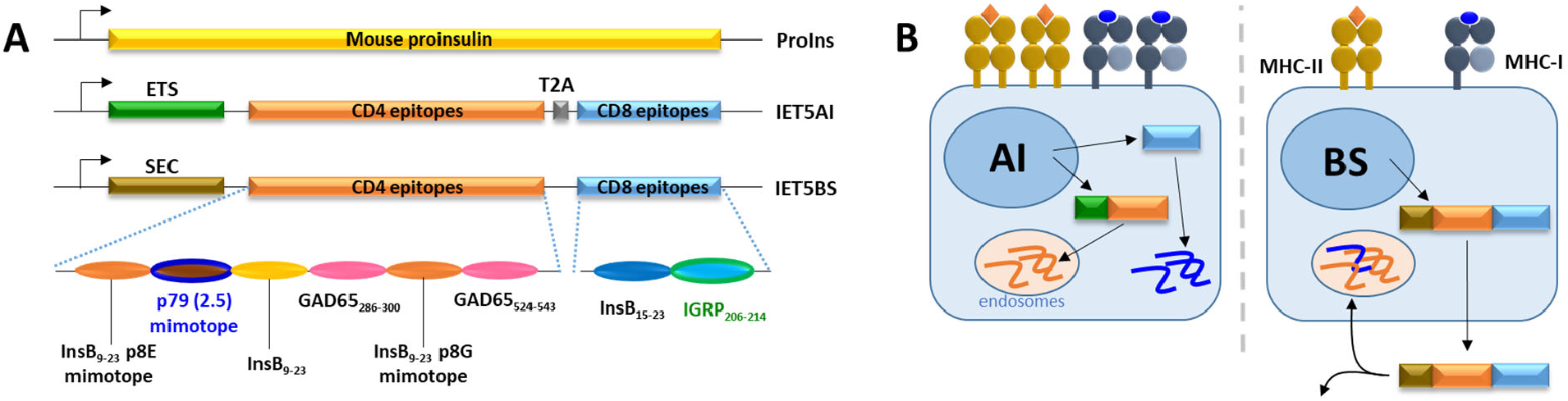
DNA constructs: arrangement and targeting of epitopes. (**A**) Constructs used for expression of mouse proinsulin (ProIns) or tailored epitopes (IET5AI and BS) on pDNA vectors (pBHT568 backbone). The AI variant features the invariant chain’s endosomal targeting signal (ETS) Ii_1-80_, preceding native CD4 epitopes (InsB_9-23_; GAD65_286-300_; GAD65_534-543_) and mimotopes (InsB_9-23_ R22E [p8E] and E21G/R22E [p8G]; and 2.5mi [p79]), followed by a cleavage site (T2A), separating them from native CD8 epitopes (InsB_15-23_; IGRP_206-214_). The BS variant features the same epitopes and mimotopes as AI but expressed them on a single polypeptide preceded by an albumin secretion signal (SEC). The epitopes recognized by the TCR-tg T cells from BDC2.5 and NY8.3 mice are indicated in blue and green, respectively. (**B**) Epitopes produced by the AI variant remain intracellular, resulting in maximal antigen loading in the transfected cells, whereas epitopes produced by the BS variant are secreted. The secreted polypeptides may then be taken up by the producing cells and by other cells in the vicinity.

**Fig. 2.**
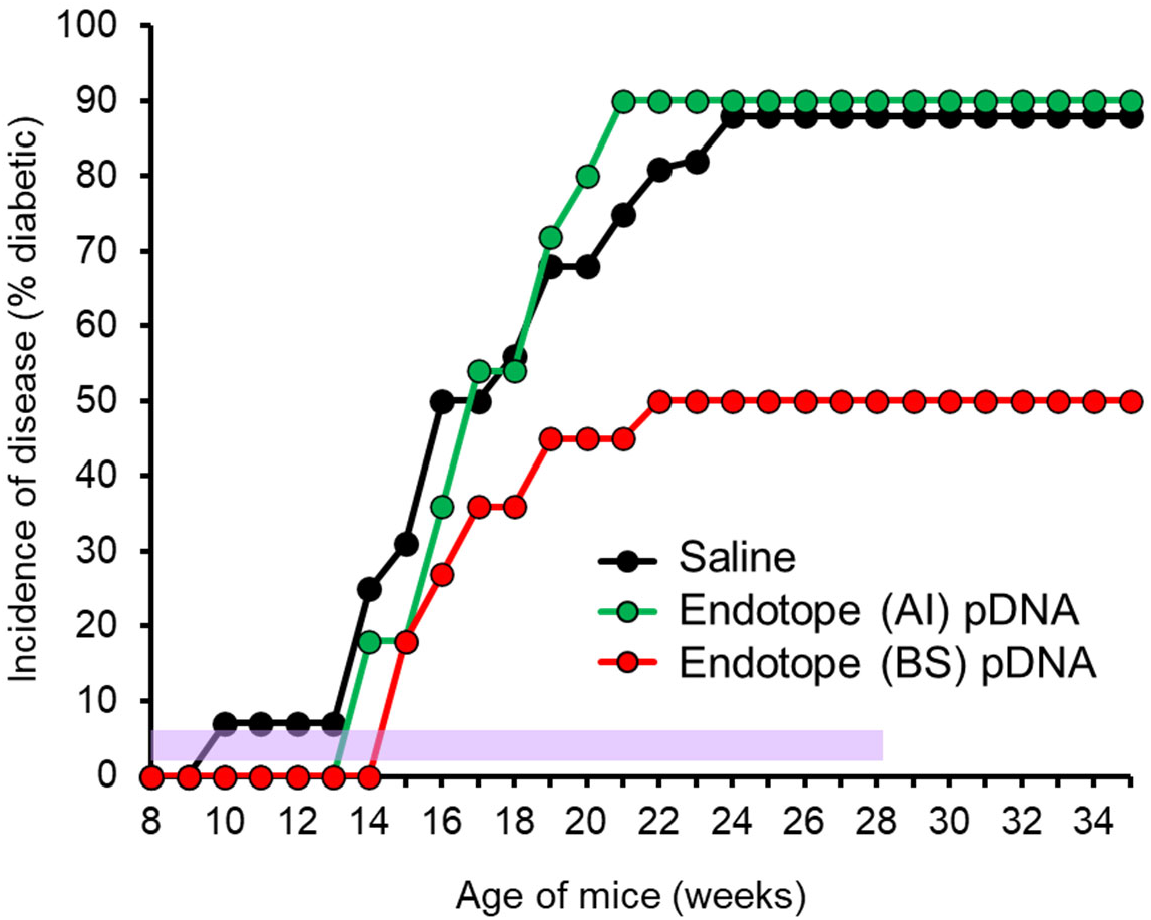
Secreted, but not intracellular, pDNA-encoded Endotope polypeptide reduced disease incidence in NOD mice. Female NOD mice (n=16 mice for saline, 11 for AI and 10 for BS) were treated i.d. weekly for 20 weeks, starting at week 8 of age (period of treatment indicated with purple shading). Log Rank test: saline vs BS: p=0.079.

### Endotope pDNA shows similar efficacy to Proinsulin pDNA with the intradermal route conferring the best protection

We next compared our selected BS Endotope pDNA with the Proinsulin pDNA, applied in a continuous weekly treatment. Using i.m. administration, both treatments reduced the incidence of disease, although it did not reach significance in this instance with the number of mice used (**Fig.3A**). Using i.d. administration, the reduced incidence of disease was significant for Endotope and Proinsulin, both at the time of treatment discontinuation and one month after (**Fig.3B**). At least during that last month without treatment, the mice appeared protected somewhat durably, without precipitous increase in onset in remaining mice.

**Fig. 3.**
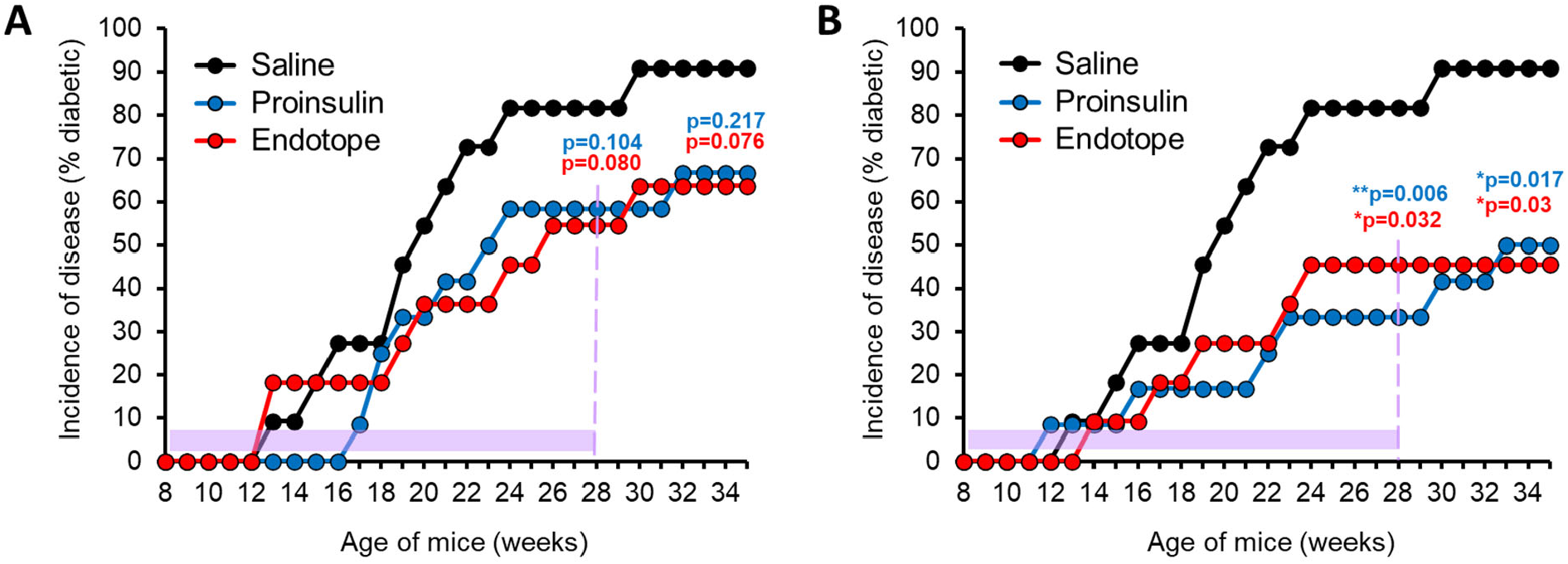
Comparable efficacy of Proinsulin and Endotope DNA vaccines, using two routes of administration. Female NOD mice (n=11 mice for saline, 12 for Proinsulin pDNA i.m. and i.d. and 11 for Endotope pDNA i.m. and i.d.) were treated i.m. (**A**) or i.d. (**B**) for 20 weeks starting at 8 weeks of age (period of treatment indicated with purple shading). Log Rank test: the p values (against saline control) at the end of treatment (28 weeks of age) and at the end of follow up (35 weeks of age) are indicated on the graph.

### Protection is lost after discontinuation of shorter treatments

To determine whether stable control of disease could be achieved with fewer treatments, mice were treated weekly for 8 weeks with Endotope or Proinsulin pDNA, either i.m. (**Fig.4A**) or i.d. (**Fig.4B**). In contrast to the previous long-term treatments, most mice rapidly developed T1D soon after treatment discontinuation in both groups and with both routes, indicating that longer continuous DNA delivery is necessary to achieve stable, long-lasting protection. Nonetheless, Endotope pDNA, but not Proinsulin pDNA, achieved significant delay in disease incidence with both routes, which was consistent with our previous report using the AI/BS mix administered over 8 weeks (23).

**Fig. 4.**
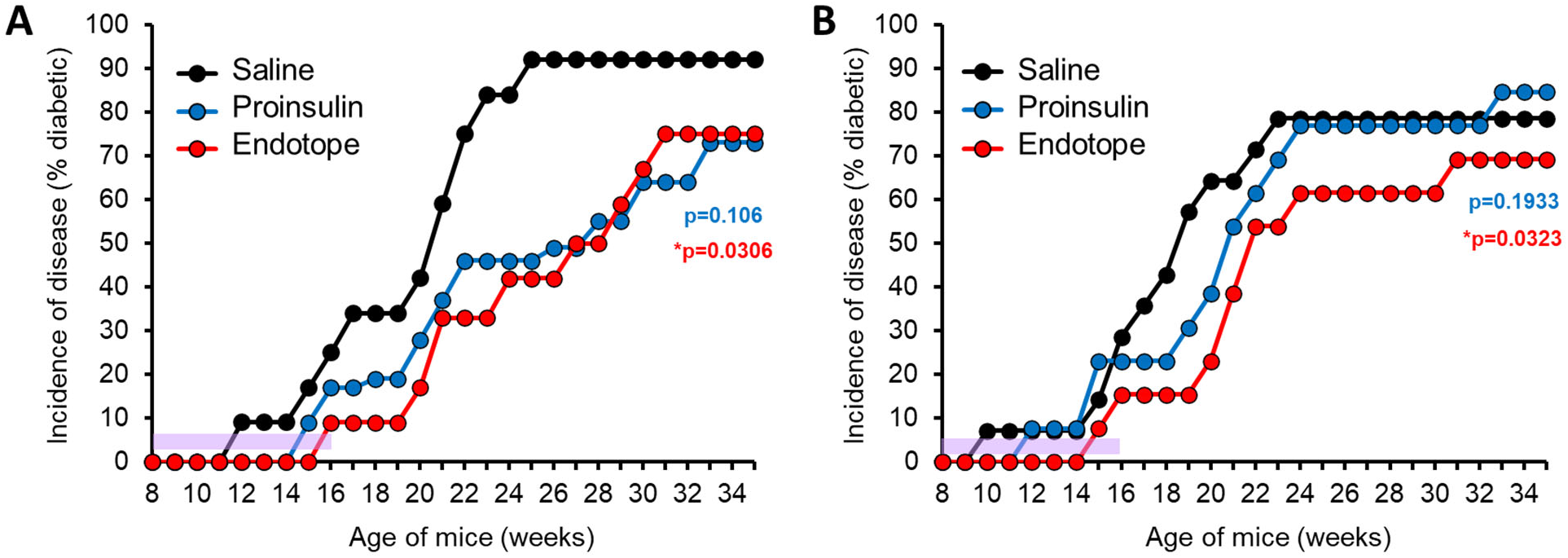
Loss of durable protection from disease following treatment discontinuation. Female NOD mice were treated i.m. (**A**) or i.d. (**B**) for 8 weeks starting at 8 weeks of age (period of treatment indicated with purple shading). Number of mice in i.m. treatment cohort: n=12 mice per group; and in i.d. treatment cohort: n=14 for saline, 13 for Proinsulin, and 13 for Endotope. Log Rank test: the p values (against saline control) at the end of follow up (35 weeks of age) are indicated on the graph.

### Persistence of antigen presentation is not informative for treatment frequency requirements

We previously demonstrated that expression of the pDNA-encoded gene in vivo, after either i.m. or i.d. delivery, lasts at least one week based on luciferase signal measurement. However, we wondered whether antigen presentation can persist even longer. To assess how long antigen remains presented after pDNA administration, we used antigen-specific CD4+ and CD8+ T cells from BDC2.5 and NY8.3 mice, respectively, which respond to two epitopes expressed by the Endotope pDNA (**Fig.1A**), labeled them with a proliferation dye and injected them into the i.d.-treated NOD mice at several time points (**Fig.5A**). The proliferative response of the T cells was analyzed 3 days after transfer. Day 0 shows the background proliferation of the T cells in absence of treatment (no proliferation in lymph nodes except in the pancreatic lymph nodes where the antigen is naturally present, draining from the islets) (**Fig.5B,C**). Proliferation of the T cells in the inguinal lymph nodes (draining the sites of pDNA inoculation) at all time points indicated that antigen was still presented as late as 14 days after treatment (**Fig.5B,C**). The response did not decrease over the two-week period for CD8+ T cells and decreased slightly (yet remain significantly above background) for the CD4+ T cells.

**Fig. 5.**
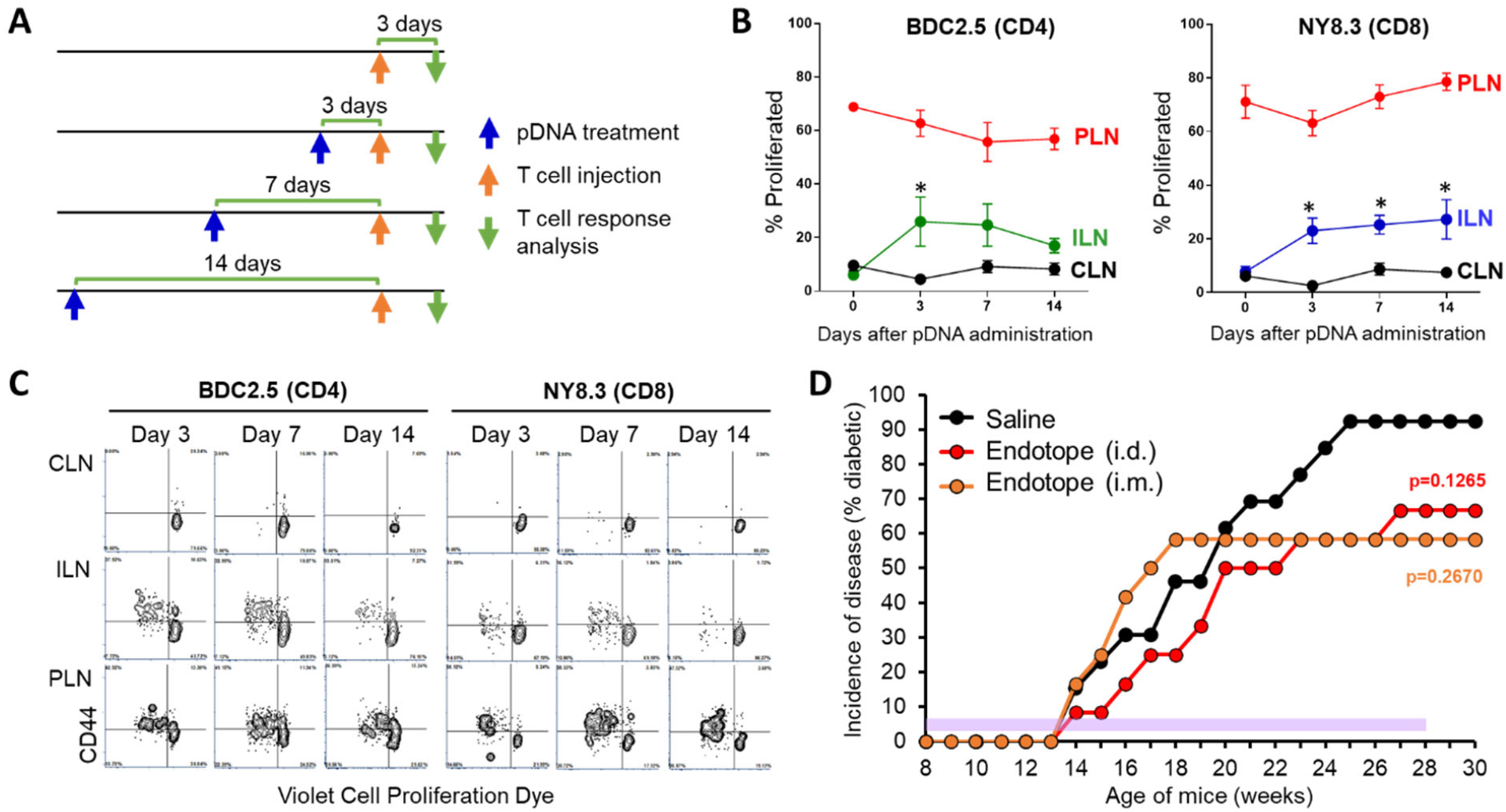
Persistence of Endotope’s epitopes presentation in vivo after pDNA inoculation and effect of reducing the frequency of treatment on disease incidence. (**A**) Four groups of female NOD mice (n=5 mice per group; except non-immunized control, n=2) were used as recipients of TCR-tg T cells from BDC2.5 and NY8.3 mice: they were treated by i.d. injection as indicated by the blue arrows; T cells were adoptively transferred as indicated by the orange arrows and 3 days after transfer, lymph nodes (LNs) from these mice were analyzed for antigen-specific T cell responses. (**B**) Proliferation of BDC2.5 CD4+ and NY8.3 CD8+ T cells (based on % T cells having divided at least once) in inguinal LNs (ILN) draining the inoculation site, cervical LNs (CLN) serving as negative control and pancreatic LNs (PLN) serving as positive control. Data show the mean ±SEM. Stars indicate significance of difference between CLN and ILN at each time point (multiple T-tests comparison). In addition, two-way ANOVA with Sidak correction was applied across the time points, indicating significant difference between CLN and ILN for BDC2.5 CD4+ T cells (**p=0.002) and NY8.3 CD8+ T cells (**p=0.0015). (**C**) Representative dot plots illustrating the data shown in panel B. (**D**) Incidence of disease in mice treated biweekly with i.m. or i.d. injections of Endotope pDNA. Female NOD mice (n= 13 mice for saline, 12 for Endotope pDNA i.d. and 12 for Endotope pDNA i.m.) were treated every other week for 20 weeks starting at 8 weeks of age (period of treatment indicated with purple shading). Log Rank test: the p values (against saline control) at the end of follow up (30 weeks of age) are indicated on the graph.

In light of these results, we postulated that although continuous treatment was important to achieve significant protection, weekly injections may be too frequent if antigen presentation persists at least two weeks. Thus, we treated NOD mice i.m. and i.d. for 20 weeks, this time every other week. Under these conditions, Endotope pDNA offered only partial and non-significant protection from T1D (**Fig.5D**), suggesting that late presentation of antigens (beyond one week) does not contribute to disease protection.

### Proinsulin pDNA provides better protection when administered at a stage immediately preceding the onset of diabetes (dysglycemic stage)

Solvason et al. previously reported that Proinsulin pDNA vaccination was particularly effective at blocking the onset of disease when NOD mice start to develop dysglycemia (150-250 mg/dL range), on their way to hyperglycemia. We tested whether the Endotope pDNA had the same capacity. When mice reached the 150-200 mg/dL range, they were treated twice a week for 6 weeks. After this treatment period, 78% of saline-treated mice had developed diabetes (glycemia > 250 mg/dL), as compared to 50% for Endotope group and 22% for the Proinsulin group (**Fig.6A**). We continued to follow these mice for another 14 weeks after the treatment ended (**Fig.6B**). Mice treated with Endotope pDNA that had not developed diabetes remained protected long-term, whereas mice treated with Proinsulin pDNA started to develop diabetes after the treatment ended to ultimately reach the same level of protection as Endotope. While diabetes onset typically occurs after 12 weeks of age, dysglycemia may be detected anywhere between 8 and 20 weeks of age. In contrast to stage 1 prevention studies, mice in this experimental set-up for stage 2 prevention are not all treated at the same age. Mice that progress to dysglycemia later may have a milder form of disease that may be easier to treat. Thus, we stratified each group of mice in two subsets: the half that developed dysglycemia the earliest (before 12 weeks approximately; **Fig.6C**) and the other half that developed dysglycemia the latest (after 12 weeks; **Fig.6D**). Interestingly, the protective effect of Proinsulin pDNA was significantly more pronounced in the second half group, suggesting that this vaccine is particularly effective at blocking the milder form of disease, while Endotope pDNA had a modest effect in both aggressive and mild settings.

**Fig. 6.**
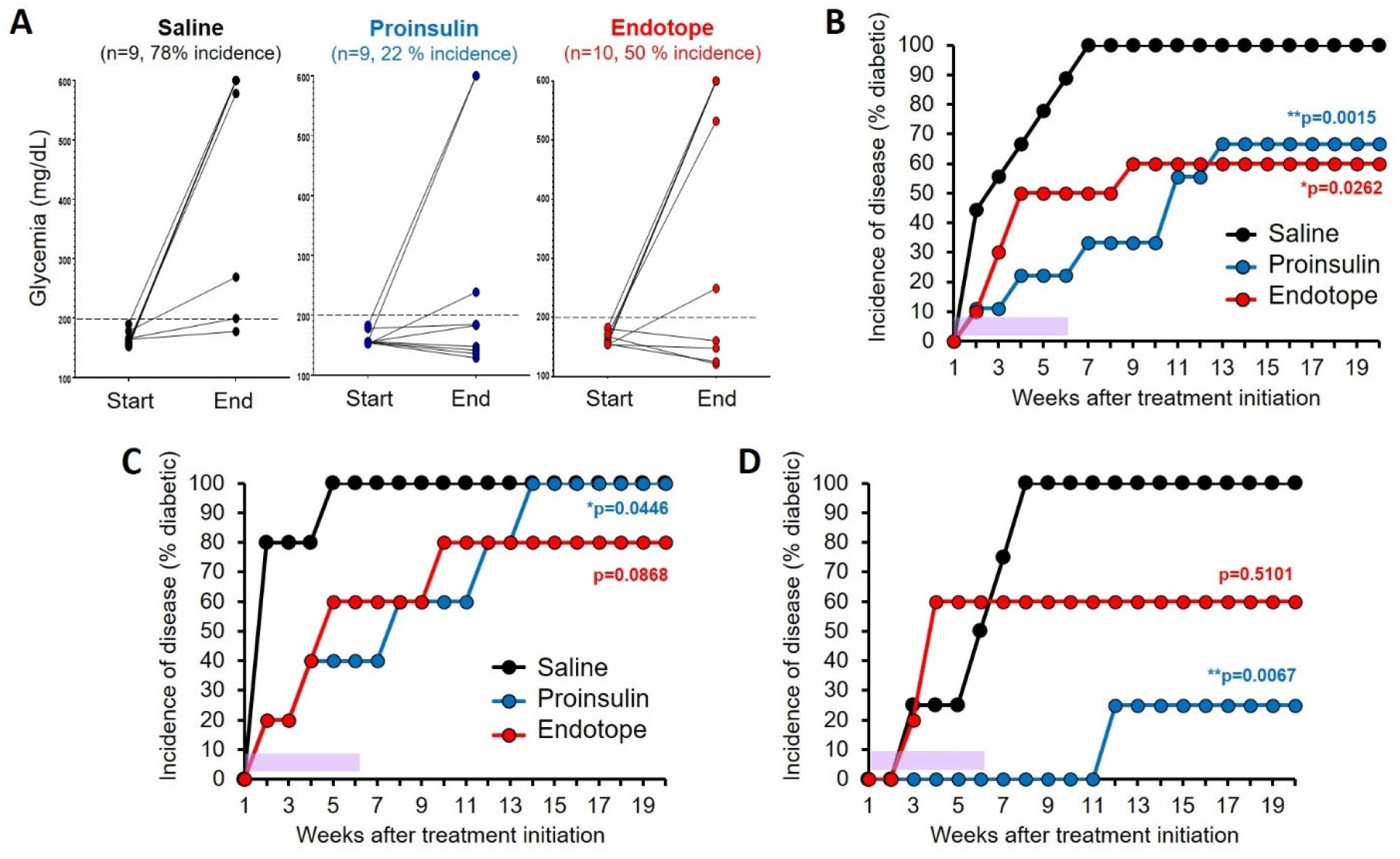
Efficacy of Endotope pDNA compared to Proinsulin pDNA in preventing progression of disease during the dysglycemic stage. Female NOD mice (n= 9 mice for saline, 9 for Proinsulin pDNA and 10 for Endotope pDNA) were enrolled when glycemia reached the 150-200 mg/dL range and treated i.d. for 6 weeks (period of treatment indicated with purple shading), with two injections per week, totaling 12 doses. (**A**) Blood glucose of mice at the start (first dose) and end (last dose) of the treatment. (**B-D**) Incidence of disease of the whole cohort (**B**), of the mice that developed dysglycemia before 12 weeks of age (n=5 mice per group) (**C**) and of the mice that developed dysglycemia after 12 weeks of age (n=4 mice for saline and Proinsulin, 5 mice for Endotope) (**D**). Log Rank test: the p values (against saline control) at the end of follow up (20 weeks of age) are indicated on the graph. The p value for Proinsulin pDNA comparing subgroups in panels C and D is p=0.0218.

## DISCUSSION

Although the inbred NOD mouse model is genetically homogeneous and permissive to the disease, there are gender-based, environmental and stochastic factors that affect the rate of spontaneous occurrence of disease in these mice (26-28) and responsiveness to treatment. Female NOD mice develop diabetes at a variable rate, between the age of 12 and 25 weeks, while males are more protected. Mice that developed dysglycemia later were significantly more responsive to Proinsulin pDNA treatment, and also to other treatments (29). In humans, individuals who develop 2+ autoantibodies at a younger age progress to diabetes the fastest (30). In addition to the genetic heterogeneity that characterizes T1D patients, diversity in immune profile (predominant autoantibodies, immunoprevalent circulating autoreactive T cells) contribute to the establishment of disease endotypes, which can help stratify patients based on shared features (12). Responsiveness to ASIT is likely linked to this immune profile. For example, responsiveness to oral insulin in the DPT-1 study was limited to patients with highest levels of insulin autoantibodies (13, 14). Accordingly, patients recruited for ASIT trials are increasingly selected based on the presence of at least the autoantibody corresponding to the administrated antigen (e.g. GAD65 in the DiAPREV-IT2 trial, NCT02387164). More clinical studies and retrospective analyses are required to determine what responders have in common and how the choice of antigens, delivery modalities and routes should be made for these subpopulations in subsequent treatments (1, 8).

Given the complexity of the human disease, it is yet unclear what fraction of the T1D patient population will benefit from the Proinsulin DNA vaccine, and the current Phase 2 SUNRISE trial (NCT03895437) is expected to provide further insights into the mechanisms of action and the factors influencing responsiveness. Proinsulin is certainly an excellent antigen choice for the NOD mouse, given its key role in disease initiation (31, 32). However, studies by two independent labs showed that Proinsulin pDNA alone (i.m. route) had a significant effect when given at stage 2 but did not reach significance when given at stage 1 of disease (24, 33). Our data mirrored those earlier observations, as we achieved more protection in stage 2 with Proinsulin pDNA after short twice-weekly treatment (22%) than in stage 1 after a long weekly treatment period (∼40%) using the i.d. route.

The Endotope platform shifts the focus from a single antigen to selected epitopes that can encompass multiple antigens and to which T cell reactivity is well documented. Important disease-driving epitopes now include various neoepitopes (6, 7, 9, 34), which can be incorporated in the Endotope design. Mimotopes can also be used to mimic certain PTMs (e.g. deamidation) or force binding on MHC in a particular register that are better recognized by autoreactive T cells (35). The fact that these epitopes are endogenously produced by the patient’s own cells from Endotope constructs also increases the chance for these peptides to be endowed with relevant PTMs. When translated to patients, this approach will require a more detailed analysis of the immune profile and disease-relevant immunopeptidome, an area that is still in its infancy despite recent progress in profiling patients with comprehensive combinatorial MHC multimer panels (19-21, 36). It will also require selection of patients based on HLA haplotypes to ensure proper presentation of the encoded epitopes, as done in trials involving the injection of peptide mixtures (NCT02620332) or peptide-pulsed dendritic cells (NTR5542) (37). To its advantage, the full protein can produce peptides that can collectively cover a wide variety of HLA haplotypes, but delivery of more than one protein in a single plasmid is technically challenging. Because another major advantage of DNA vaccination is the ability to design and manufacture plasmids cheaply and in large quantities, it is an attractive modality when it comes to producing multiple versions for groups of patients. Thus, Endotope constructs would be particularly suitable for precision medicine of T1D using ASIT.

The present study serves as preclinical proof-of-concept that a selection of relevant epitopes from various antigens recognized by dominant T cell clones may be as effective as a single disease-driving protein antigen (Proinsulin) in blocking or delaying the progression of autoimmune diabetes. In all studies involving treatments during late stage 1 disease (all mice treated at the same age while normoglycemic), Endotope and Proinsulin pDNA performed similarly, with the i.d. route appearing more suitable, at least for NOD mice. In short treatments, however, Endotope treatment delayed the onset of disease more than Proinsulin. In contrast, when mice were treated upon entering the dysglycemic phase (stage 2), a difference between the two vaccines was more evident. The effect of Endotope was the same whether the mice entered stage 2 early or late, whereas as discussed above, Proinsulin had a major impact on those entering stage 2 late. The advantage of Proinsulin in late stage 2 disease may be potentially related to broader epitope coverage within Proinsulin if intramolecular epitope spreading occurs primarily during the peri-insulitis to insulitis transition. It is unclear how many of the Endotope-encoded epitopes significantly contribute to the protection achieved. Rivas et al. showed that expression of just one of the BDC2.5 mimotopes (also targeted toward the MHC-II processing pathway in a different way) was sufficient to significantly reduce the incidence of disease (17). More recently, we showed that two peptides in combination (p79 mimotope and hybrid insulin peptide (2.5HIP)) protected 70% of mice using a different delivery modality (soluble antigen array) (5). Treatment with a single InsB9-23 (R22E) mimotope (p8E, also present in our Endotope construct) in soluble form also significantly prevented the onset of disease in NOD mice (38). Presentation of a few selected epitopes on MHC molecules displayed on nanoparticles in Pere Santamaria’s successful studies also leads to tolerance that spread to other epitopes of the same antigen or others (16, 39). Collectively, these studies illustrate the fact that ASITs are progressively shifting their focus from antigens to epitopes. This may be particularly important in the case of hybrid peptides, which cannot be found in a single protein. Moreover, it is possible that hybrid peptides and PTM peptides are produced exclusively in the periphery under specific conditions and may not be found in the thymus to mediate negative selection. Thus, special consideration should be given to these neoepitopes in ASITs (7). The Endotope constructs from these studies should prove useful to the field as they contain epitopes recognized by T cells from many strains of TCR transgenic NOD mice that are widely used (BDC2.5, NY8.3, BDC12-4.1, G9C8).

An important insight from these studies is the observation that the secreted version of the polypeptide containing all epitopes has therapeutic benefit, but not the version in which peptides are targeted intracellularly. On the one hand, this is reminiscent of another DNA vaccine study in which the long GAD65_190-315_ peptide was more protective when secreted than the full non-secreted GAD65 protein in NOD mice (40). On the other hand, we have shown that both secreted and non-secreted forms induced immune responses in draining lymph nodes (23). Thus, poor efficacy was not a result of a lack of antigen presentation, although T cell responses tended to be lower in amplitude with the non-secreted version (23). It is possible that among the APCs involved, those not directly transfected and relying on antigen uptake play a more important role in tolerance induction. However, despite not being secreted or not being targeted to endosomes, which should in both cases result in better presentation of MHC class II-restricted epitopes, the Proinsulin vaccine was comparable to Endotope vaccine in efficacy. Thus, it would be interesting to test whether a secreted Proinsulin vaccine would have a more pronounced effect in treated animals, in addition to other optimization strategies (41). Interestingly, expression of preproinsulin, which can more efficiently localize to the endoplasmic reticulum, can lead to unwanted CD8+ T cell responses that can exacerbate disease, demonstrating that careful subcellular targeting of antigens can make a significant difference between immunogenic and tolerogenic response (42).

Finally, how long should delivered antigens be expressed to achieve durable tolerance is a critical question to which there is still no clear answer. At least in the NOD mouse model, the disease can be significantly delayed as long as mice are under regular treatments, but disease progression resumes soon after cessation of treatment. It is expected that the maintenance of anergy and the continuous deletion of new autoreactive T cells requires uninterrupted exposure to antigen presented under tolerogenic conditions (to offset the presentation of the same antigens in inflamed pancreatic lymph nodes). In contrast, induction of regulatory T cells (Tregs) should confer long-lasting protection providing that these Tregs are functionally stable, which may not be the case in both NOD mice and T1D patients (43, 44). We have previously shown that presentation of CD4 epitopes from Endotope pDNA does result in a significant increase in antigen-specific Foxp3+ Tregs (23). Induction of Tregs appears to be consistent with the fact that expression of a few epitopes was sufficient to broadly control diabetogenic T cells and disease progression as a result, whereas anergy/deletion should only impact the T cells that are specific to the delivered antigens. However, our data also support the idea that these induced Tregs do not persist and continuous exposure to autoantigens, possibly away from the pancreatic lymph nodes, is therefore needed to control autoreactive T cells. As transfected pDNA can reside in cells in episomal form, DNA vaccines appear ideal to accomplish prolonged antigen exposure. We therefore addressed how long would antigens be presented after a single injection (two epitopes, one presented on MHC-II and the other on MHC-I, were tested). Despite significant stimulation of antigen-specific T cells in lymph nodes draining the vaccination site being measured at least two weeks later, reducing the frequency of inoculations from once a week to once every two weeks resulted in substantial reduction in therapeutic efficacy. This surprising finding underlines the limitation of basing treatment ASIT frequency on the persistence of antigen presentation. It is possible that the two epitopes tested are of sufficiently high affinity that reduction in antigen levels did not lead to noticeable changes in T cell responses.

Overall, these studies provide important new insights on the use of tolerogenic pDNA vaccines to treat autoimmunity, including proof-of-principle that expression of selected epitopes tailored to groups of patients (or a specific strain of mouse in this case) can achieve protection comparable to that a full antigen known to drive the disease. However, issues of Treg stability and antigen persistence remain major hurdles to overcome for the successful treatment of T1D.

## MATERIALS AND METHODS

### Plasmid constructs

All constructs used in these studies were previously described (23) and expressed in the pBHT568 (originally containing Proinsulin) from Lawrence Steinman and Peggy Ho (24). The epitopes expressed (**Fig.1A**) have all been well described to engage diabetogenic T cells in NOD mice (22, 23). Construct A intracellular (AI) contains the endosome targeting signal (Ii_1-80_) selected from a previous study (22) and a T2A cleavage site. This construct generates two polypeptides that are differentially targeted to MHC class II or MHC class I (**Fig.1B**) (22). Construct B secreted (BS) contains the albumin secretion signal, and all epitopes are secreted as one polypeptide to be disseminated to other antigen-presenting cells (**Fig.1B**) (23).

### Mice

All mouse strains were purchased from The Jackson Laboratory and bred in our barrier facility: NOD (#001976), NOD.CD45.2 (#014149) and T cell receptor transgenic (TCR-Tg) mice BDC2.5 (#004460) and NY8.3 (#005868). TCR-Tg T cells from BDC2.5 and NY8.3 mice respectively recognize the p79/2.5 mimotope (2.5mi) presented on MHC class II (I-A^g7^) and the IGRP_206-214_ epitope presented in MHC class I (K^d^), all encoded by our constructs (**Fig.1A**). Male and female mice donors and recipients were used at 8-16 weeks of age for studies of T cell response and female NOD mice were used at 10 weeks of age for preclinical experiment unless otherwise indicated. All studies were approved by Columbia University’s Institutional Animal Care and Use Committee.

### Disease prevention studies (Stage 1)

NOD females (10 weeks of age) were treated by either intramuscular (i.m.) or intradermal (i.d.) injection, weekly or biweekly for a duration of 8 weeks or 20 weeks as indicated in the figures. Each dose consisted of 50 μg of pDNA, encoding either Proinsulin or an Endotope construct, in 100 μL of saline. Control groups, used to determine the normal incidence of disease, received 100 μL of saline. At the time of treatment, mice were normoglycemic but are not known to have peri-insulitis and autoantibodies at this age (26); this corresponds to Stage 1 disease in humans (45). The i.m. injection was split between the two quadriceps, while i.d. administration was split between the two flanks of the abdominal area after shaving. Mice from different groups were mixed in cages to minimize cage variability. Blood glucose was monitored weekly (up to 35 weeks of age) using Prodigy glucometer and test strips. Mice were diagnosed as diabetic after two consecutive blood glucose levels greater than 250 mg/dL and sacrificed upon diagnosis or at the end of the observation period if normoglycemic.

### Antigen persistence study

NOD female mice (8-10 weeks of age), used as recipients, were immunized i.d. with 50 μg of Endotope-BS pDNA at days −14, −7 and −3 before adoptive transfer of TCR-Tg T cells from BDC2.5 and NY8.3 donor female mice (8-16 weeks of age). Spleen and pooled lymph nodes were collected from donor CD45.2+ BDC2.5 and NY8.3 mice, and CD4+ CD25- and CD8+ T cells were purified using the MojoSort Mouse CD4 (supplemented with biotinylated anti-CD25) and CD8 T Cell Isolation Kits (BioLegend), respectively. Cells were then labelled with Violet Cell Proliferation Dye (eBioscience) and 0.5-1×10^6^ T cells were injected i.v. into recipient NOD (CD45.1+) mice to all groups on day 0. Antigen-specific congenic CD4+ and CD8+ T cells were analyzed by flow cytometry (BD Fortessa) 3 days later. Data were analyzed with FCS Express 7.

### Disease stabilization studies (Stage 2)

NOD female mice were monitored for blood glucose levels from 10 weeks of age and were considered dysglycemic when in the range of 150-200 mg/dL (this corresponds to Stage 2 disease in humans (45)) and subsequently enrolled in the vaccination study, following 2 doses weekly for a period of 6 weeks (12 doses total), groups were randomly assigned on a rotating basis to intradermal treatment with saline, Proinsulin or Endotope pDNA until all animals were enrolled. Mice were diagnosed as diabetic after two consecutive blood glucose levels greater than 250 mg/dL and sacrificed when reaching > 500 mg/dL over two consecutive readings or at the end of the observation period if normoglycemic.

### Statistical analysis

All statistical testing was performed using GraphPad Prism 4.0. For diabetes incidence, Log Rank (Mantel-Cox) test was used. Two-way ANOVA with Sidak correction and multiple T-test comparisons were performed in other studies as indicated in legends.

## Data availability

All study data are included in the article.

## ACKNOWLEDGMENTS

JPF was funded by a postdoctoral fellowship from the American Diabetes Association (1-18-PDF-151). The study was supported by the Translational Therapeutics Accelerator program funded by the National Center for Advancing Translational Sciences, National Institutes of Health, through Grant Number UL1TR001873. Some experiments involved the use of the Columbia Center for Translational Immunology Flow Cytometry Core, supported in part by the Diabetes Research Center (grant P30DK063608).

